# Social evaluation of skilfulness in Tonkean macaques (*Macaca tonkeana*) and brown capuchins (*Sapajus apella*)

**DOI:** 10.1101/2025.05.13.653791

**Authors:** Marie Hirel, Michele Marziliano, Hélène Meunier, Hannes Rakoczy, Julia Fischer, Stefanie Keupp

## Abstract

For optimal decision-making, social animals can benefit from evaluating others’ behaviours. Some species seemingly consider the skills of others when deciding who to interact with in different contexts. Yet, whether and how nonhuman animals form impressions about others’ competence is still unclear. In this study, we investigated whether Tonkean macaques (*Macaca tonkeana*) and brown capuchins (*Sapajus apella*) can evaluate the skilfulness of others. Subjects observed two human actors (one skilful, one unskilled) trying to open several food containers. Only the skilful actor successfully opened the containers and released food so the experimenter could give it to the subjects. Afterwards, subjects did not choose the skilful actor significantly more frequently than the unskilled one. Their choices for the skilful actor did not increase through trials, nor were they based on the outcomes experienced in previous trials. However, when we considered their initial preferences for the human actors, we observed a significant shift in preference for the skilful actor. Our subjects also looked preferentially at the skilful over the unskilled actor when both simultaneously manipulated a container. We observed substantial individual variation and species differences in choice and looking behaviours. While the cognitive mechanisms (impression formation vs outcome-based process) underlying these social decisions are still unclear, our findings indicate that Tonkean macaques and brown capuchins used social information about the actors’ skills to inform their decisions.

## Introduction

Forming impressions about others from their past behaviours is a valuable cognitive skill, for example, in social contexts that require choosing interaction partners or role models (Corriveau et al., 2009; Fusaro et al., 2011; Hermes et al., 2016, 2017; Kushnir et al., 2013; Rakoczy et al., 2009; Wu et al., 2016). Most research has focused on evaluating others’ prosocial characteristics in nonhuman primates (Anderson et al., 2016; Anderson, Kuroshima, et al., 2013; Anderson, Takimoto, et al., 2013; Herrmann et al., 2013; Kawai et al., 2014; Russell et al., 2008; Subiaul et al., 2008) and a few other animal species (dogs and wolves: Jim et al., 2022; Kundey et al., 2011; fishes: Bshary & Grutter, 2006; Vail et al., 2014; elephants: Jim et al., 2021; cats: Leete et al., 2020). Social evaluation of others’ skills (*i.e.*, the performance of an individual in a particular task; Sih et al., 2019) is also essential social information to consider when deciding who to interact with. Individuals might benefit from observing and/or interacting with the skilful foragers of their group to access food resources or to learn new foraging skills. Similarly, recruiting the most skilful partners seems advantageous for successful cooperation.

Some field experiments show that several nonhuman primates increased their affiliative behaviours toward the only group member skilled at providing food in a novel experimental foraging situation (Fruteau et al., 2009; O’Hearn et al., 2025; Stammbach, 1988). Others preferentially observed (Ottoni et al., 2005; Tan et al., 2018) or copied (Canteloup et al., 2021; Horner et al., 2010; Kendal et al., 2015) the most skilful group members at a foraging technique. However, only a few experimental studies investigated the ability to evaluate the skills of others in nonhuman species. After observing two human actors repeatedly trying to open containers filled with food, long-tailed macaques (*Macaca fascicularis*) preferentially chose (Placì et al., 2019), and dogs preferentially looked at or approached (Chijiiwa et al., 2022; Piotti et al., 2017) a skilful over an unskilled actor. Chimpanzees (*Pan troglodytes*) and coral trout (*Plectropomus leopardus*) strategically recruited the most skillful conspecific partners for cooperation (Keupp & Herrmann, 2024; Melis et al., 2006; Vail et al., 2014).

These previous studies suggest that some animal species can consider the skills of others and behave differently towards individuals who differ in their degree of skilfulness. Yet, whether nonhuman species adapt their behaviour toward skilful individuals because they have formed impressions about their skills or because of other cognitive processes is still unclear. For example, confounding factors such as individual differences other than skills, familiarity, or pre-existing relationships between the subjects and the individuals being assessed were not always considered. To broaden our understanding of social evaluation in nonhuman primates, we investigated whether two monkey species, Tonkean macaques (*Macaca tonkeana*) and brown capuchins (*Sapajus apella*), can evaluate the skilfulness of others at making food accessible.

Both species live in multi-male multi-female societies characterised as tolerant (De Waal, 2000; Fragaszy et al., 2004; Riley, 2010; Thierry et al., 1994, 2004; Thierry, 2007). They demonstrated social cognitive skills such as social relationships knowledge, attention and goal-directed understanding, and visual perspective-taking (Canteloup et al., 2016, 2017; Canteloup & Meunier, 2017; Defolie et al., 2015; Fragaszy et al., 2004; Phillips et al., 2009; Whitehouse & Meunier, 2020). They exhibit tool-using in foraging contexts (Anderson, 1985; Ducoing & Thierry, 2005; Fragaszy et al., 2004; Ottoni & Izar, 2008), and brown capuchins even showed preferential attention toward skilful conspecific foragers (Ottoni et al., 2005). Tonkean macaques had never been tested on social evaluation before (except in Hirel et al., 2024) while brown capuchins already demonstrated social evaluation of human actors’ helpfulness and reciprocity (Anderson, Kuroshima, et al., 2013; Anderson, Takimoto, et al., 2013). Therefore, we hypothesised that both species could evaluate others’ skills.

In this experiment, subjects could sample information about two human actors’ skills from observation during two demonstration sessions. One actor (skilful) always succeeded at opening transparent containers filled with food, while the other actor (unskilled) always failed. To assess the animals’ evaluation of the actors, we then measured the proportion of choosing the skilful actor and of anticipatory behaviours (*i.e.*, time looking at or being close to the actors).

In the Choice Test, subjects could choose between the two actors for eight trials. If subjects chose the skilful actor, they received a food reward from the experimenter once the container was successfully opened by the actor. If they chose the unskilled actor, they went empty-handed. We predicted that 1) both species would choose the skilful actor more often than the unskilled actor, and 2) shift their initial actor’s preference before the experimental manipulation in favour of the skilful actor as this became the better option. We also examined whether the subjects, rather than sampling and evaluating the actors’ skills, engaged in a Win-stay/Lose-shift (WS/LS) strategy by basing their decisions on the outcome of the previous test trial (like the chimpanzees in Melis et al., 2006). In the Expectation Test, we measured subjects’ anticipatory behaviours while the two actors simultaneously manipulated a food container for 20 s. Preferential looking or spatial proximity towards the skilful actor would indicate anticipation of the successful task completion and that the subjects expected the outcome of the actors’ actions. We predicted that the subjects’ anticipatory behaviours would increase toward the skilful actor while decreasing toward the unskilled actor as subjects obtained more and more information about the actors’ skills.

## Methods

### 1 Subjects and study site

This study was conducted at the Centre de Primatologie – Silabe de l’Université de Strasbourg (France) between July and September 2023. Nineteen subjects participated in the study including ten Tonkean macaques (four females; 13.7 ± 5.0 years old) from two different social groups and nine brown capuchins from one social group (five females; 8.3 ± 4.4 years old; Table S1), all living in wooded outdoor enclosures with constant access to indoor rooms (more details on subjects and housing can be found in the Supplementary Materials). Individuals were fed once per day with dry pellets and once per week with fruits and vegetables, and had access to water *ad libitum*. Subjects were tested individually in experimental rooms situated next to their outdoor enclosure (Figure S1). They participated in the experiment on a voluntary basis, which may have biased our sample of subjects towards dominants, individuals who are more comfortable in the experimental rooms and/or experienced in experimental testing (see STRANGE framework; Webster & Rutz, 2020). The subjects had participated in several ethological studies, but they had never been tested with social evaluation experiments before (except for one Tonkean macaque in Hirel et al., 2024; Table S1). One brown capuchin only completed the ExpT1 and three trials of the Choice Test but was included in the analyses as specified below.

### 2 Materials

Two humans, both being well familiar to the subjects, played the roles of the skilful and the unskilled actors. The task was to open transparent containers filled with food. The skilful actor was always successful at opening the containers to release the food, whereas the unskilled actor always failed. The actors’ roles were constant for each subject throughout the experimental procedure and were attributed according to subjects’ choices in the Initial Preference Assessment (see below). Differently looking containers (in colours of screw caps and in sizes: small: 120 mL, medium: 180 mL, large: 250 mL) were used to emphasise the actors’ (in)ability to open any kind of screw-top containers. Both actors manipulated the same combination of size and coloured screw caps during the demonstration sessions while only the large containers with the yellow screw cap were used for the other experimental steps. Containers were filled with raisins for about ^2^/_3_ of the volume and raisins were used as rewards. At each step, the actors sat at two predetermined locations in front of the mesh of the testing room, 1.5 m apart and with a distance from the mesh adjusted for the arm-length of each subject. The actors carried their containers in identical baskets and, once opened, emptied them into transparent plastic bowls (750 mL). Two identical wooden sticks (length = 30 cm) ending with yellow wooden squares (side = 8 cm) were used as targets.

### 3 Procedure and design

The experiment consisted of eight steps divided into three ‘working’ sessions for each subject (Figure 1a and Table 1). Each subject underwent up to one working session per half day. To increase the possibility for the subjects to evaluate the actors’ skills at manipulating the containers, the subjects experienced the possible states of the containers (*i.e.*, open or close) and the outcomes in their food intake (Kuroshima et al., 2014), and how a human can open and close the containers, during a familiarisation session before the test phase. In addition, as touching the target was the measure of subjects’ choices for the actors during the IPA and Choice Test, the experimenter ensured that each subject was able to choose the better of two options by touching targets, before the experiment. Only subjects who met the learning criterion (*i.e.*, choosing the better food option on at least ten out of 12 trials in a session) could participate in the study (for more details about the familiarisation and target check, please refer to the Supplementary Materials).

**Figure 1:**
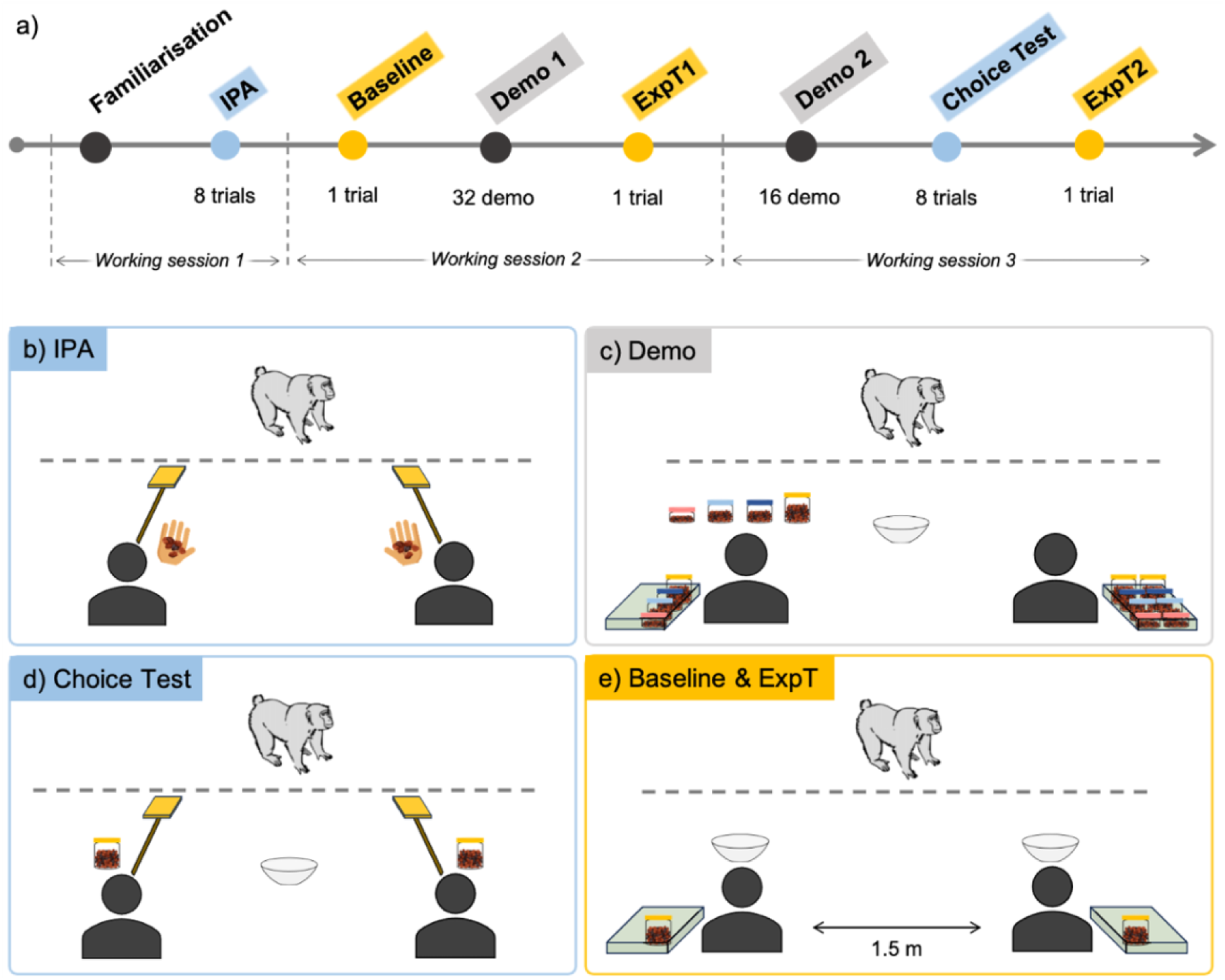
a) Chronological order of the experimental steps; b) Materials’ and actors’ disposition at the start of each trial of the Initial Preference Assessment (IPA): the two actors presented a target in one hand and raisins on their palm of their other hand; c) Demonstration sessions: the two actors sat in front of the mesh with their basket and containers next to them, four containers in front of the active demonstrator, the bowl in the middle (note that the actors have eight containers each to manipulate in Demo 1 and four in Demo 2); d) Choice Test: the two actors sat in front of the mesh and presented their target, one container was placed in front of each actor, the bowl was in the middle; e) Expectation Test (Baseline, ExpT1, ExpT2): the two actors sat in front of the mesh with their basket and one container next to them, a bowl was placed in front of each actor, each actor manipulated one container. For each step, the actors sat at the same two predetermined locations in front of the mesh of the testing room, 1.5 m apart and with a distance from the mesh adjusted for the length of each subject.

**Table 1:** Information about the different experimental steps

| Step | Type | Measure | Purpose | Analyses |
| --- | --- | --- | --- | --- |
| Familiarisation | 1 session | - | Let subjects experience themselves the possible states of the containers and how they can be opened and closed | - |
| Demonstration | Demo 1 (8 obs / actor) Demo 2 (4 obs / actor) | Looking time | Give subjects the opportunity to obtain information about the actors' skills from observation | - |
| Initial Preference Assessment (IPA) | 1 session (8 trials) | Actor choices | Assessing subjects' initial preferences for the actors and assigning the roles of the actors | Estimation of the change in preference for the skilful actor from the IPA to the Choice Test |
| Choice Test | 1 session (8 trials) | Actor choices | Measuring the number of choices of the subjects for the skilful actor | Estimation of the probability to choose the skilful actor according to trial number |
| Expectation Test | 3 trials of 20 s each: Baseline, ExpT1, ExpT2 | Looking & spatial proximity time | Measuring the subjects' time of looking at and being close to the actors before the demonstrations (Baseline), after Demo 1 (ExpT1), and after both demonstrations and Choice Test (ExpT2) | Estimation of the effect of trial (Baseline, ExpT1, ExpT2) on the subjects' time of looking at and/or in proximity to both actors |

#### 3.1 Initial preference assessment (IPA)

We conducted eight IPA trials before the experimental manipulation to assess whether subjects had a spontaneous initial preference for one of the actors, *i.e.*, a personal preference unrelated to any experimental manipulation. First, the two actors fed the subject a few raisins, one after the other, to show the subject that both were willing to give food. Then, each IPA trial started with both actors moving simultaneously to their designated location, holding one raisin in the palm of their hand and a target in the other hand. Once the experimenter lured the subject to the central position equidistant to the actors, the actors presented their target simultaneously in front of them, and the experimenter stepped back. The subject could approach one of the actors to touch their target and obtain the food; this target touch was considered a choice for this actor (video S1). The actors’ locations were pseudo-randomised (*i.e.*, an equal number of times on each side) within subjects and counterbalanced between subjects. If the subjects chose each actor in four out of eight trials, the actors’ roles were randomly assigned and counterbalanced between subjects. If a subject chose one actor five times or more out of the eight trials, we assigned to this ‘preferred’ person the role of the unskilled actor for this subject. We chose this conservative role assignment rule to avoid a confound of initial preference for an actor and choice based on skill demonstrations of the actor.

#### 3.2 Baseline

We measured the baseline for subjects’ anticipatory behaviours before they acquired any information about the skilfulness of the actors during the demonstration sessions. The two actors moved simultaneously to their designated location and placed their basket containing one identical closed baited container next to them. Two transparent bowls, previously placed by the experimenter, were in front of them, out of reach for the subject (Figure 1e). Once the experimenter lured the subject to the central position equidistant to the actors, the actors started simultaneously manipulating their container while the experimenter stepped back (video S5). After 20 seconds of manipulation, the skilful actor successfully opened their container and emptied it into their bowl while the unskilled actor tried vainly to empty their still closed container into their bowl. Both actors then simultaneously put their containers into their baskets and looked down. The experimenter first took the empty bowl in front of the unskilled actor to show it to the subject, then took the full bowl in front of the skilful actor to feed the subject with a few raisins. During the duration of the trial, the actors focused only on the containers and never looked at the subject.

#### 3.3 Demonstration sessions

Each demonstration session began with the two actors moving simultaneously to their designated location, holding a target and placing their basket containing identical sets of baited containers next to them. One transparent bowl was in the middle equidistant from the actors, out of reach for the subject (Figure 1c). Each actor alternately demonstrated their skill to open four containers in a row in one location (subsequently referred to as one demo trial). In the Demo 1 session, each actor had eight closed baited containers in their basket and alternately demonstrated twice in the same location. The actors then repeated the same procedure but at the opposite location. In the Demo 2 session, the procedure was the same except that the actors held an identical set of four containers and only one demo trial per location was performed by each actor. Therefore, the subjects observed each actor manipulating 16 containers (eight containers per side) during Demo 1 and eight containers (four containers per side) during Demo 2, resulting in a total of 24 observations per actor for each subject. The actors’ order and location were pseudo-randomised and counterbalanced between and within subjects.

A demo trial started with an actor placing four containers aligned in front of them, out of reach for the subject, and then presenting their target. In the meantime, the other actor stayed head down without moving until the end of the demo trial. Once the subject had touched the target, the actor started manipulating the four containers. After trying for around five seconds, the skilful actor successfully opened each container, emptied it into the transparent bowl, put the container and the lid back into their basket, and repeated the same actions with the next three containers. The unskilled actor did the same actions, except that they attempted to open the container for around five seconds but failed, and tried vainly to empty the closed container into the transparent bowl. We are aware that five seconds may appear relatively brief for an unsuccessful attempt; however, it was important to keep the manipulation time equal between successful and unsuccessful attempts, to avoid confounding a potential preference/avoidance for one of the roles with this actor merely being associated with longer handling of the container. As the monkeys’ attention span is relatively short, we opted for the compromise of five seconds to avoid introducing too much additional noise in the data due to a loss of interest by the subjects. After manipulating their fourth container, the actor stopped moving and looked down. The experimenter took the transparent bowl which was either filled with food (after a skilful actor’s manipulation) or empty (after an unskilled actor’s manipulation). The experimenter then fed the subject some of the food from the bowl or showed the empty bowl to the subject, before replacing the bowl with a new identical empty one (video S2). We deemed it necessary to reward the animals with small quantities of food, but not directly from the skilful actor; otherwise, they might have lost the motivation to participate in the experiment.

#### 3.4 Choice Test

The subjects chose in eight trials which of the two actors could manipulate a container to release food. A trial began with the two actors moving simultaneously to their designated location, placing one identical closed baited container in front of them and holding a target in their hand. One transparent bowl was in the central position, equidistant to the actors, out of reach for the subject. Once the experimenter lured the subject to the central position, the actors presented their target simultaneously while the experimenter stepped back (Figure 1d). The subjects indicated their choice by approaching and touching the target of one actor. The unchosen actor removed their target, put their container back, and waited with their head down until the end of the trial. Once chosen, the skilful and unskilled actors followed the same actions with their container as during the demonstration trials. The experimenter then fed the subject some of the food from the bowl (after the skilful actor’s manipulation) or showed the empty bowl to the subject (after the unskilled actor’s manipulation), before replacing the bowl with a new identical empty one (videos S3 and S4). The subjects were given a maximum of 20 seconds to make a choice, otherwise the trial was aborted. A trial (with the same actors’ location configuration) was repeated a maximum of three times before aborting the session (which never happened). The actors’ location was pseudo-randomised within subjects and counterbalanced between subjects.

#### 3.5 Expectation Test

We measured subjects’ anticipatory behaviours toward the actors in two trials, after subjects had information about the actors’ skills from Demo 1 (ExpT1) and from both demonstrations and the Choice Test (ExpT2; Figure 1a). The procedure for ExpT1 and ExpT2 was exactly the same as for the Baseline (videos S5 and S6). The actors’ location was the same for the three trials for one subject, but counterbalanced between subjects.

### 4 Data coding

All experimental steps were videotaped with three GoPro9 cameras to obtain different views of the subjects, the actors, setups and experimental rooms. All the videos were coded frame by frame by an observer using Behavioral Observation Research Interactive Software (BORIS v.8.20; Friard & Gamba, 2016). A second observer, who was unaware of the study design and hypothesis, coded independently 38 videos which were pseudo-randomly selected to include 20% of the sessions for each combination of phase, species and subjects, giving a relatively representative sample of each behaviour coded. Inter-coder reliability scores ranged between 63% to 100% (see supplementary materials for details). For the IPA and Choice Test, videos were coded for: a) the choices of the subjects toward either actor, b) the latency of choices for the subjects, c) the synchronicity of targets’ presentation by the actors, and d) the amount of time subjects spent in close range to either actor (1 m radius), in nearby range or far away (Figures S1 and S2). For the Expectation Test and the demonstration sessions, videos were coded for: a) the duration of looking at either actor, b) the position of the subjects in the experimental room (close, nearby, or far away). The exact definitions of the coded behaviours are reported in Table S2.

### 5 Data analyses

#### 5.1 Choices

We assessed whether subjects chose the skilful actor more often than the unskilled actor in the Choice Test and whether this probability of choosing the skilful actor increased with direct experience (*i.e.*, trial number), by fitting a Generalized Linear Mixed model (GLMM; Baayen, 2008) with logit link function (McCullagh & Nelder, 1989) and a binomial error structure. The sample included 147 observations (*i.e.*, trials) from 19 subjects (including the brown capuchin subject who completed only the first three trials of the Choice Test session). The response variable for each trial was 1 (choice of the skilful actor) or 0 (choice of the unskilled actor). The fixed-effect predictors we included in this model were trial number (main predictor), species (capuchin, Tonkean), the attention directed at the actors during the demonstration, and the three-way interaction between these predictors. This three-way interaction was included to account for the possibility that the two species, but also the subjects who were more or less attentive during the demonstration could have different trial learning curves, or that the two species may require different amounts of time to acquire comparable levels of information about the actors’ skills. As a measure of the subjects’ attention to both actors, we chose the minimum looking time per subject to the skilful and unskilled actor, *i.e.*, the time during which each subject observed each actor. To control for their potential effects, we included the sex of the subjects, the side location (left, right), and the identity (A, B) of the skilful actor. An additional control predictor, synchronicity of targets’ presentation (-1: unskilled actor first, 0: synchronous, +1: skilful actor first), was added to the model. During video coding, we noticed that the two actors were not always fully synchronous which could have affected the choices of the subjects.

To account for individual differences, avoid overconfident model estimates, and keep type I error rate at the nominal level of 5%, we included subject ID as a random intercept effect and all identifiable random slopes within subject, which were trial number, side location of the skilful actor, and synchronicity of targets’ presentation (Barr et al., 2013; Schielzeth & Forstmeier, 2009). As an overall test of the fixed effects and to avoid cryptic multiple testing (Forstmeier & Schielzeth, 2011), we compared this full model with a null model lacking the effect of trial number and its interactions but being otherwise identical, using likelihood ratio tests (Dobson, 2018). To test the impact of individual fixed effects, we conducted likelihood ratio tests (Dobson, 2018) that compared the full models with reduced models, each lacking one fixed effect at a time (Barr et al., 2013). We obtained confidence intervals of model estimates and fitted values using a parametric bootstrap (N = 1000 bootstraps). We checked all the relevant model assumptions and transformed some variables when needed to ease the interpretation of the model estimates and convergence. This procedure of assumption checks applies to all analyses (see supplementary materials for more details on each model analysis).

#### 5.2 Shift in preference for the skilful actor

Regardless of whether the probability of choosing the skilful actor during the Choice Test differed from chance (50%), subjects might have chosen the skilful actor more often in the Choice Test than during the IPA. Therefore, we fitted a similar model except that we estimated the change in preference for the skilful actor from the IPA to the Choice Test. The sample included 288 observations (*i.e.*, trials) from 18 subjects (excluding the brown capuchin subject who did not complete the Choice Test). The response variable was a matrix comprising the number of choices for the skilful and unskilled actors for each subject and phase. This model included the test phase (IPA, Choice Test) as the main predictor and species (capuchin, Tonkean). To account for random individual differences, avoid overconfident model estimates, and keep the type I error rate at the nominal level of 5%, we included subject ID as a random intercept effect and the only theoretically identifiable random slope phase within the subject.

#### 5.3 Alternative WS/LS strategy

The subjects could have used alternative strategies rather than sampling and evaluating the actors’ skills. Like chimpanzees in the study by Melis et al. (2006), they could engage in a WS/LS strategy by tracking their success with each of the actors and basing their decisions on the outcome of the previous trial. We ran a post-hoc GLMM analysis to estimate the effect of positive outcomes on the probability of staying with or switching actor choices during the Choice Test. For each trial, we coded whether the subjects stayed with or switched their choice of actors from the previous trial (variable ‘strategy’) and whether they had chosen the skilful actor and obtained a food reward in the previous trial (variable ‘reward’). The sample included seven trials per subject (trials 2 to 8), resulting in 126 trials in total. In the model, we included strategy (stay, switch) as the response variable, reward (yes, no) as a fixed effect, and species as a control predictor. We also included subject ID as a random intercept effect to account for random individual differences, to avoid overconfident model estimates and to keep the type I error rate at the nominal level of 5%. If subjects follow the WS/LS strategy, they should base their decisions on the outcomes in the previous trial. We would observe more “stay” decisions after successes than failures and more “switch” decisions after failures than successes.

#### 5.4 Anticipatory behaviours

We assessed whether subjects would preferentially look or be close to the skilful actor while both actors simultaneously manipulated a baited container for 20 s. We wanted to estimate the effect of trial (Baseline, ExpT1 and ExpT2) on the subjects’ looking time and their proximity to both actors, as subjects gained more indirect and direct information about the actors’ skills. As the response variables were bound between 0 and 20 s, we turned them into proportions to rule out fitted values and confidence intervals of fitted values extending beyond the possible response range. We fitted two GLMMs with a beta error distribution (McCullagh & Nelder, 1989) and with identical structure except for the response variable, which was either the proportion of time looking at or the proportion of time spent in proximity to either actor. The sample for each model included 112 observations from 19 subjects and 56 levels of trial nested in subject (including the brown capuchin subject who completed only the Baseline and ExpT2). In these models, we included actor (skilful, unskilled), trial (Baseline, ExpT1, ExpT2) and their interaction as the main predictors. To control for their potential effects, we also included species, the sex of the subjects, and the identity of the skilful actor.

To account for random individual differences, to avoid overconfident model estimates and to keep the type I error rate at the nominal level of 5%, we included subject ID as a random intercept effect and all theoretically identifiable random slopes (Barr et al., 2013; Schielzeth & Forstmeier, 2009), which were trial and actor within subject. In addition, we included the random intercept effect of trial nested in subject to account for the fact that the data for the skilful and unskilled actor for any given combination of subject and trial were not independent. Following the same model comparison procedure as before, we compared these full models with null models lacking the effects of trial, actor and their interaction but being otherwise identical, and with reduced models, each lacking one fixed effect at a time, using likelihood ratio tests.

#### 5.5 R functions and packages

We conducted statistical analyses and created plots using R (version 4.3.2; R Core Team, 2022). We used the function glmer of the package lme4 (version 1.1-35.1; Bates et al., 2015) to fit the logistic models, and the function glmmTMB of the homonymous package (version 1.1.8; Brooks et al., 2017) for the models with beta distribution. Parametric bootstraps were obtained using the functions bootMer (package lme4) and simulate (package glmmTMB).

## Results

### 1. Choices

In the Choice Test, subjects chose the skilful actor 85 times out of 144 (brown capuchins: 39 choices, 60.9%; Tonkean macaques: 46 choices, 57.5%) and they chose the unskilled actor 59 times out of 144 (brown capuchins: 25 choices, 39%; Tonkean macaques: 34 choices, 42.5%; excluding the brown capuchin who did not complete the session). Twelve subjects (six out of nine brown capuchins and six out of ten Tonkean macaques) chose the skilful actor at the first trial (Table S1). Tonkean macaques chose the skilful actor 57.5% of the time in the first four trials and same in the last four trials, while brown capuchins chose 46.9% the skilful actor in the first four trials and 75% in the last four trials. Overall, their probability of choosing the skilful actor in the Choice Test was not influenced by any of the predictor terms however (full-null model comparison: χ^2^ = 7.345, df = 4, *p* = 0.119; Table 2). This result indicates that subjects did not choose the skilful actor more often than expected by chance, nor did they increase their choices for the skilful actor over the trials.

**Table 2:**
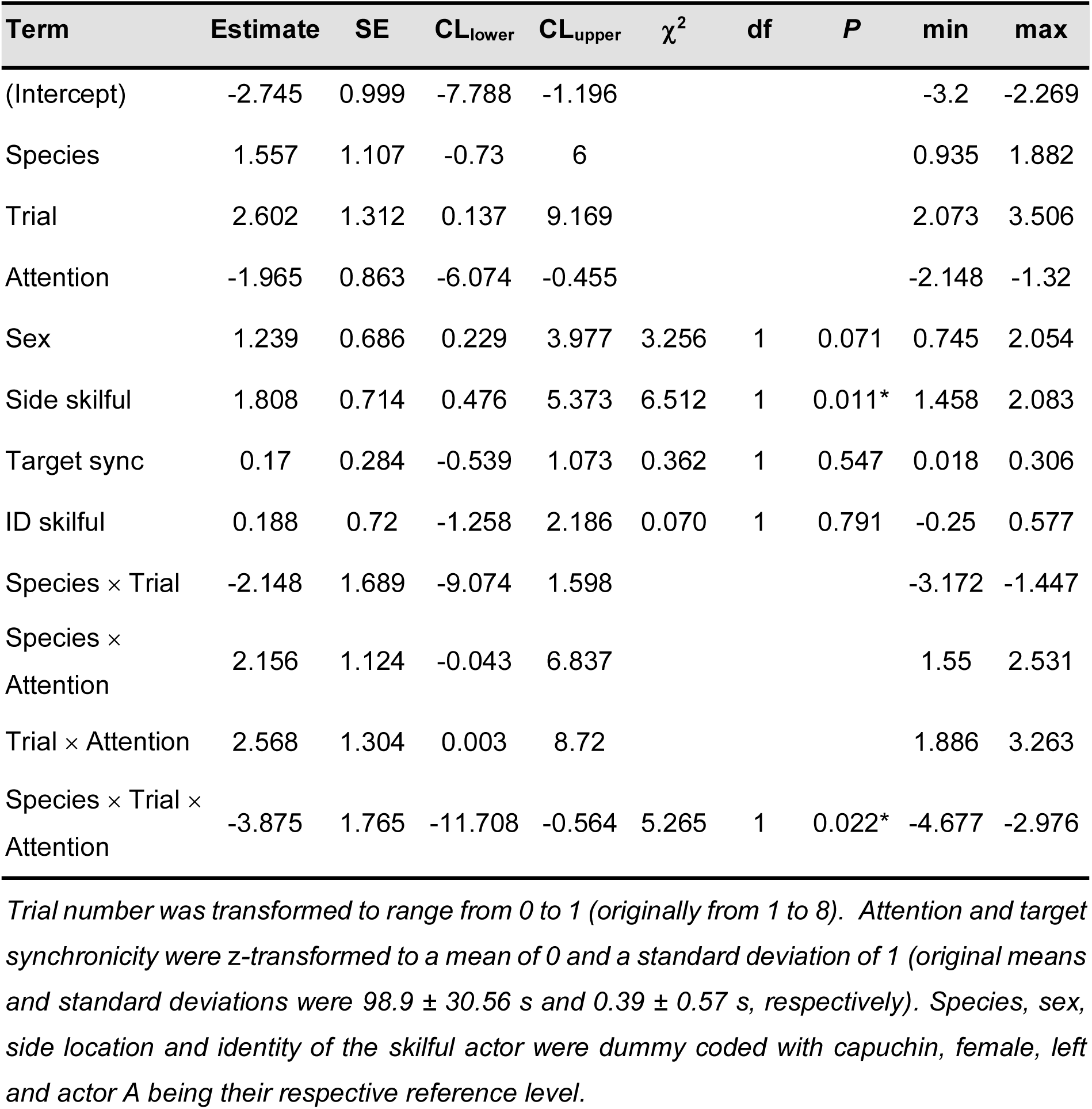
Results of the subjects’ choices full model in the Choice Test (estimates together with standard errors, 95% confidence limits, significance tests and the estimates range obtained when dropping levels of grouping factors one at a time).

| Term | Estimate | SE | CL<sub>lower</sub> | CL<sub>upper</sub> | $\chi^2$ | df | P | min | max |
| --- | --- | --- | --- | --- | --- | --- | --- | --- | --- |
| (Intercept) | -2.745 | 0.999 | -7.788 | -1.196 |  |  |  | -3.2 | -2.269 |
| Species | 1.557 | 1.107 | -0.73 | 6 |  |  |  | 0.935 | 1.882 |
| Trial | 2.602 | 1.312 | 0.137 | 9.169 |  |  |  | 2.073 | 3.506 |
| Attention | -1.965 | 0.863 | -6.074 | -0.455 |  |  |  | -2.148 | -1.32 |
| Sex | 1.239 | 0.686 | 0.229 | 3.977 | 3.256 | 1 | 0.071 | 0.745 | 2.054 |
| Side skilful | 1.808 | 0.714 | 0.476 | 5.373 | 6.512 | 1 | 0.011* | 1.458 | 2.083 |
| Target sync | 0.17 | 0.284 | -0.539 | 1.073 | 0.362 | 1 | 0.547 | 0.018 | 0.306 |
| ID skilful | 0.188 | 0.72 | -1.258 | 2.186 | 0.070 | 1 | 0.791 | -0.25 | 0.577 |
| Species × Trial | -2.148 | 1.689 | -9.074 | 1.598 |  |  |  | -3.172 | -1.447 |
| Species × Attention | 2.156 | 1.124 | -0.043 | 6.837 |  |  |  | 1.55 | 2.531 |
| Trial × Attention | 2.568 | 1.304 | 0.003 | 8.72 |  |  |  | 1.886 | 3.263 |
| Species × Trial × Attention | -3.875 | 1.765 | -11.708 | -0.564 | 5.265 | 1 | 0.022* | -4.677 | -2.976 |

*Trial number was transformed to range from 0 to 1 (originally from 1 to 8). Attention and target synchronicity were z-transformed to a mean of 0 and a standard deviation of 1 (original means and standard deviations were $98.9 \pm 30.56$ s and $0.39 \pm 0.57$ s, respectively). Species, sex, side location and identity of the skilful actor were dummy coded with capuchin, female, left and actor A being their respective reference level.*

The model revealed a significant three-way interaction between trial, species, and the subjects’ attention during the demonstration phase (*p* = 0.022; Tables 2 and S3, Figure S3), arguably due to multiple testing. There were no significant effects of sex (*p* = 0.071), synchronicity in target presentation (*p* = 0.547), or the identity of the skilful actor (*p* = 0.791; Table 3). Success probability was higher when the skilful actor was on the right-hand side (*p* = 0.011; Table 2). In the Choice Test, subjects chose the left side 34.7% of the time and the right side 65.3% of the time, and four subjects chose the right side in at least seven out of the eight trials. During the IPA, subjects chose the left side 30.9% of the time and the right side 69.1% of the time. These results indicate a general preference for the right side from the start of the experiment, which may have interfered with the subjects’ choices during the Choice Test.

**Table 3:** Results of the full model for the comparison of subjects’ choices between the IPA and the Choice Test (estimates together with standard errors, 95% confidence limits, significance tests and the estimates range obtained when dropping levels of grouping factors one at a time).

| Term | Estimate | SE | CL<sub>lower</sub> | CL<sub>upper</sub> | $\chi^2$ | df | P | min | max |
| --- | --- | --- | --- | --- | --- | --- | --- | --- | --- |
| (Intercept) | -0.648 | 0.23 | -1.162 | -0.193 |  |  |  | -0.714 | -0.571 |
| Phase | 1.021 | 0.286 | 0.482 | 1.591 | 10.228 | 1 | 0.001* | 0.891 | 1.127 |
| Species | 0.028 | 0.266 | -0.522 | 0.586 | 0.011 | 1 | 0.917 | -0.064 | 0.11 |

*The reference levels for phase and species were respectively IPA and brown capuchins.*

### 2. Shift in preference for the skilful actor

Thirteen out of the 18 subjects increased their number of choices for the skilful actor from the IPA to the Choice Test, with seven of them choosing the skilful actor in at least six out of the eight trials of the Choice Test (Table S1; Figure 2). The model revealed a strong effect of phase (*p* < 0.001; Table 3), with the probability of choosing the skilful actor increasing from 34.7% in the IPA to 59% in the Choice Test (excluding the brown capuchin’s choices who did not complete the Choice Test; Figure 2). The model revealed no species differences in this shift in actors’ preference (*p* = 0.917).

**Figure 2:**
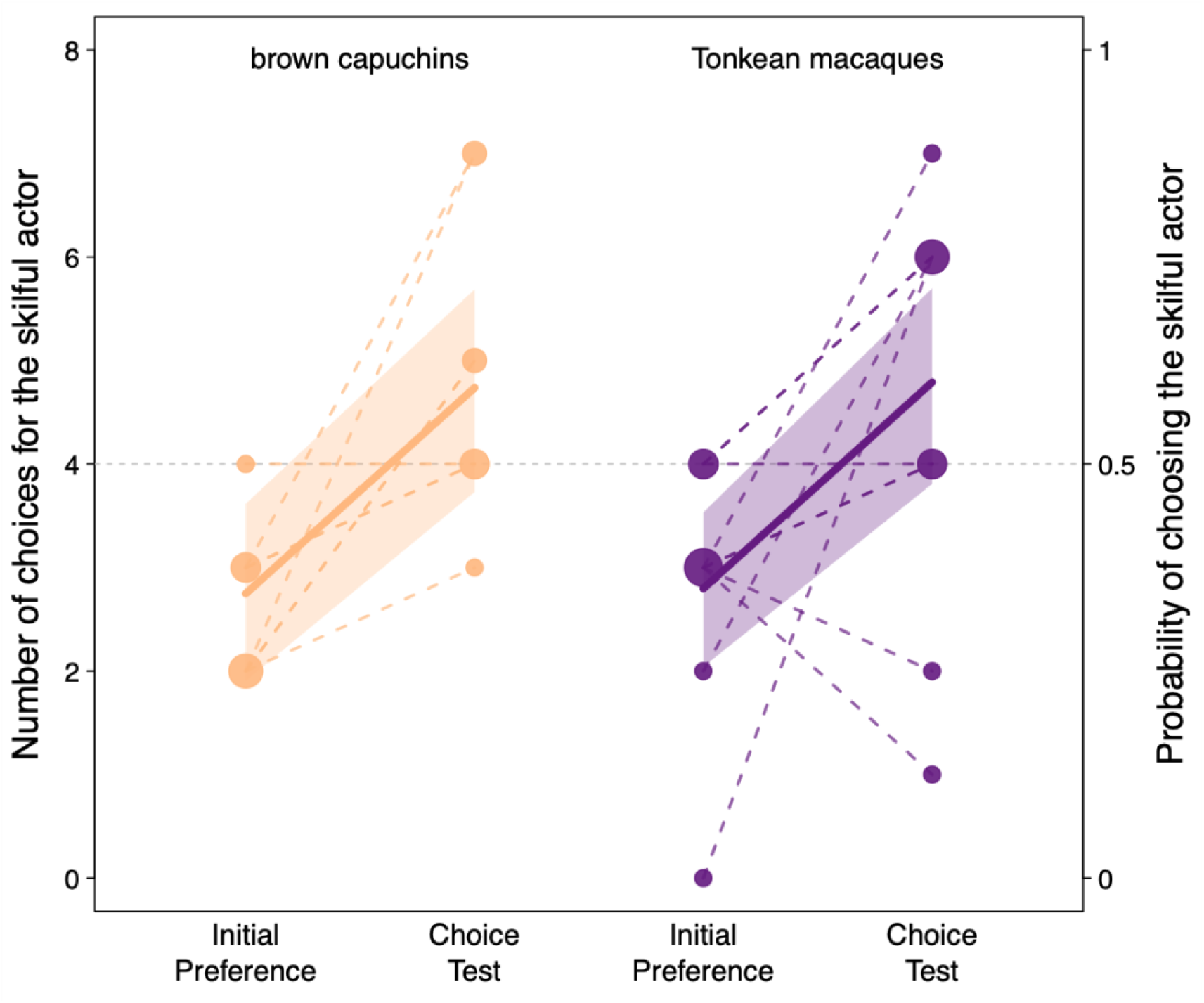
Probability of choosing the skilful actor during the Initial Preference Assessment (IPA) and the Choice Test for brown capuchins and Tonkean macaques. The continuous lines and the shaded areas depict the fitted model and its 95% confidence intervals. Points depict observations; the area of the point is proportional to the number of observations (range: 1 to 5). Observations from the same subject are connected by dashed lines.

### 3. Alternative WS/LS strategy

After an unsuccessful trial in the Choice Test (receiving no reward), the subjects stayed with the same actor choice in 23 out of 50 trials (46%) while switched in 27 out of 50 trials (54%). After a successful trial in the Choice Test (receiving a reward), the subjects stayed with the same actor choice in 46 out of 76 trials (60.5%) while switched in 30 out of 76 trials (39.5%). The model revealed a non-significant effect of reward (*p* = 0.115) and no species differences in the probability of using the WS/LS strategy (*p* = 0.709; Table 4). Even though the subjects tended to stay with the same actor more often after a win than after a loss, they did not switch more often after a loss than after a win (Figure 3). These results indicate that our subjects did not base their decisions on the outcomes they experienced in previous trials with a WS/LS strategy.

**Figure 3:**
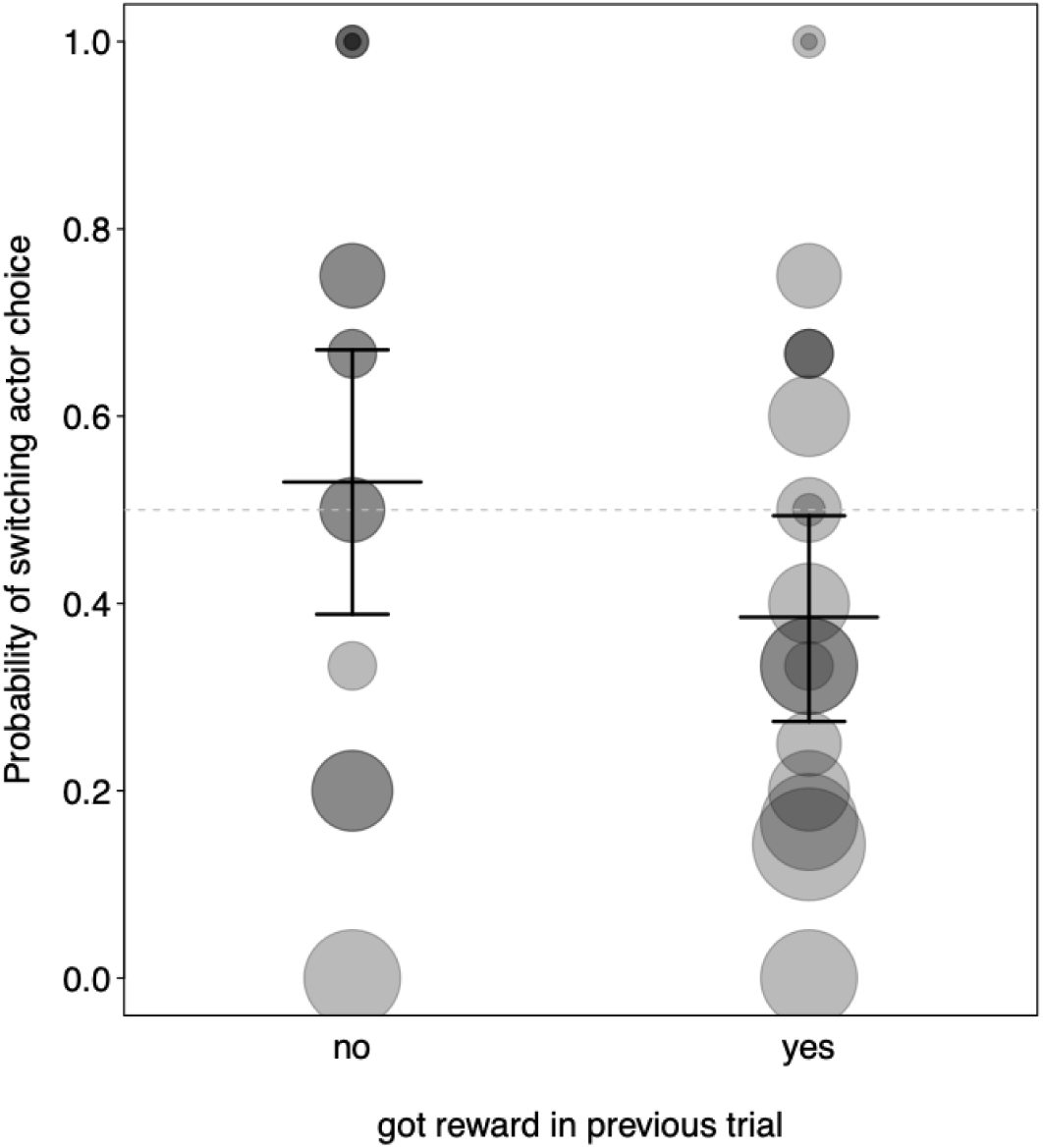
Probability of subjects switching their actor choices depending on the outcome in previous trial, *i.e.*, whether the subjects received a reward or not. Points depict observations; the area of the point is proportional to the number of observations.

**Table 4:** Results of the full model for subjects’ Win-Stay/Loose-Switch (WS/LS) strategy (estimates together with standard errors, 95% confidence limits and the estimates range).

| Term | Estimate | SE | CL<sub>lower</sub> | CL<sub>upper</sub> | $\chi^2$ | df | P | min | max |
| --- | --- | --- | --- | --- | --- | --- | --- | --- | --- |
| (Intercept) | 0.198 | 0.356 | -0.522 | 0.922 |  |  |  | 0.031 | 0.437 |
| Reward | -0.585 | 0.374 | -1.375 | 0.146 | 2.488 | 1 | 0.115 | -0.905 | -0.402 |
| Species | -0.142 | 0.380 | -0.898 | 0.686 | 0.140 | 1 | 0.709 | -0.301 | 0.041 |

*Reward was dummy coded with its reference level being no reward obtained in previous trial. The reference level for species was brown capuchins.*

### 4. Anticipatory behaviours

Overall, the proportion of time looking at the actors during the Expectation Test was significantly influenced by trial and the actors’ role (full-null model comparison: χ^2^ = 18.5, df = 5, *p* = 0.002; Table 5). During ExpT1 and ExpT2, with all other factors being averaged, subjects looked longer at the skilful (ExpT1: 4.99 s; ExpT2: 6.44 s) than the unskilled actor (ExpT1: 2.93 s; ExpT2: 2.91 s; Figure S4 and Table S4). They looked less at the unskilled actor during ExpT1 and ExpT2 than in the Baseline (5.42 s) but did not spend more time looking at the skilful actor than in the Baseline (6.52 s; Figure S4 and Table S4). The model revealed significant effects of species, with Tonkean macaques looking at the actors 60.6% longer than brown capuchins (*p* = 0.005; Tables 5 and S4) and of the skilful actor’s identity (accounting for a difference of about 1.3 s in looking time; *p* = 0.031), while the effect of sex was not significant (*p* = 0.097; Tables 5 and S4).

**Table 5:** Results of the full model for the subjects’ looking time during the Expectation Test (estimates together with standard errors, 95% confidence limits, significance tests and the estimates range obtained when dropping levels of grouping factors one at a time).

| Term | Estimate | SE | CL<sub>lower</sub> | CL<sub>upper</sub> | $\chi^2$ | df | P | min | max |
| --- | --- | --- | --- | --- | --- | --- | --- | --- | --- |
| (Intercept) | -1.072 | 0.271 | -1.638 | -0.585 |  |  |  | -1.204 | -0.968 |
| Phase ExpT1 | -0.491 | 0.255 | -1.023 | -0.008 |  |  |  | -0.595 | -0.365 |
| Phase ExpT2 | -0.106 | 0.250 | -0.595 | 0.391 |  |  |  | -0.280 | 0 |
| Actor role | -0.277 | 0.317 | -0.941 | 0.310 |  |  |  | -0.431 | -0.112 |
| Species | 0.622 | 0.197 | 0.259 | 1.048 | 7.842 | 1 | 0.005** | 0.508 | 0.821 |
| Sex | 0.329 | 0.199 | -0.050 | 0.746 | 2.757 | 1 | 0.097 | 0.172 | 0.423 |
| ID skilful | -0.452 | 0.196 | -0.841 | -0.067 | 4.660 | 1 | 0.031* | -0.614 | -0.362 |
| Phase ExpT1 × actor role | -0.309 | 0.365 | -1.001 | 0.438 | 3.007 | 2 | 0.222 | -0.448 | -0.167 |
| Phase ExpT2 × actor role | -0.650 | 0.369 | -1.362 | 0.115 |  |  | <sup>1</sup> | -0.876 | -0.442 |

*Phase, actor role, species, sex and ID of the skilful actor were dummy coded with their reference level being respectively baseline, skilful, capuchins, female and actor A.*
<sup>1</sup>only one p-value because the interaction was tested as a whole.

Overall, the proportion of time spent in proximity to the actors during the Expectation Test was not influenced by any of the predictors (full-null model comparison: χ^2^ = 2.28, df = 5, *p* = 0.808; Table S5). The proportion of time spent close to the actors was generally low and only slightly higher close to the skilful actor (ExpT1: 2.99 ± 5.44 s; ExpT2: 4.43 ± 7.47 s) than close to the unskilled actor (ExpT1: 1.38 ± 3.35 s; ExpT2: 1.29 ± 3.98 s; Table S4). However, subjects did not spend time close to each actor equally at random in the Baseline (skilful: 3.28 ± 4.78 s, unskilled: 1.44 ± 3.15 s), which may have interfered with the test results afterwards.

### Discussion

#### Shift in preference in favour of the skilful individual

We investigated whether Tonkean macaques and brown capuchins can evaluate the skilfulness of human actors at opening food containers. When choosing which of the two actors they wanted to manipulate a container to extract food, subjects’ choices for the skilful actor did not differ from chance as a group (Choice Test). However, Tonkean macaques and brown capuchins successfully shifted their initial preference for the actors in favour of the skilful actor when this switch allowed them to maximise reward outcomes in the Choice Test. Both Tonkean macaques and brown capuchins paid attention to the actors’ actions during demonstrations around half the time, suggesting they might have obtained information from observation. In addition, the absence of an increase in choices for the skilful actor through trials and a WS/LS strategy suggests that these monkeys did not base their decisions on the outcomes they experienced during the Choice test. Therefore, like other primate species (Fruteau et al., 2009; Horner et al., 2010; Kendal et al., 2015; Keupp & Herrmann, 2024; Melis et al., 2006; O’Hearn et al., 2025; Placì et al., 2019; Stammbach, 1988), Tonkean macaques and brown capuchins adapted their behaviours towards skilled individuals.

Our results indicate that our subjects paid attention to the actors’ actions and that the experimental manipulation (*i.e.*, demonstration sessions) impacted their decisions. However, we cannot disentangle whether they adapted their behaviours because they have formed impressions about the actors’ skills or because they associated the outcomes experienced with each actor (*i.e.*, outcome-based process). Indeed, subjects received small quantities of food from the experimenter after the demonstrations of the skilful actor in contrast to the demonstrations of the unskilled actor. This procedure was a compromise between not giving them any food at all – which would ensure a consistent experimental procedure but considerably reduce their level of motivation – and giving them small quantities of food not provided directly by the skilful actor, resulting in a different reward history following one type of demonstration while keeping subjects motivated to participate and pay attention. This conundrum highlights one of the challenges of investigating social evaluation abilities in nonhuman animals, as sometimes it is not possible to rely exclusively on indirect information sampling to seed the experimental manipulations (see for examples: Chijiiwa et al., 2022; Hirel et al., 2024; Keupp & Herrmann, 2024).

#### Anticipatory behaviours toward the skilful individual

Subjects looked significantly longer at the skilful actor than the unskilled actor when both actors simultaneously manipulated their containers (Expectation Test). Although they did not increase their time spent looking at the skilful actor, they significantly decreased their time looking at the unskilled actor, after the demonstrations. This preferential looking towards the skilful actor is consistent with previous findings on dogs showing a preferential looking for skilful over unskilled experimenters (Chijiiwa et al., 2022), and on brown capuchins and long-tailed macaques showing preferential observation towards skilful tool users over less skilled ones among conspecifics of their group (Ottoni et al., 2005; Tan et al., 2018). Unlike their looking pattern, subjects neither spent more time near the skilful actor than the unskilled actor nor changed their time in proximity to the actors. This result contrasts with previous findings on chimpanzees showing longer time spent in proximity to a human actor they have observed giving food to a third party before, compared to one that did not (Russell et al., 2008), and on dogs who spent more time close to skilful over unskilled experimenters (Chijiiwa et al., 2022).

Forming impressions of others’ skills could help to select the most appropriate models to learn from – a strategy maximising social learning benefits (Camacho-Alpízar & Guillette, 2023; Laland, 2004). Selective attention in social learning for competent and high-social-status group members has indeed been found in several species (*e.g.*, Canteloup et al., 2021; Horner et al., 2010; Kendal et al., 2015; Kulahci et al., 2016). For brown capuchins, the preferential attention towards skilful nutcrackers was not explained by social affiliations or social proximity between group members (Ottoni et al., 2005). In our study, food-driven attention is an unlikely explanation for the preferential looking of our subjects, as the two actors had the same amount of food in their containers, and we measured the subjects’ looking time during the 20 seconds before the skilful actor released the food. Both findings could support the hypothesis that monkeys’ looking behaviours might have been driven by previous assessments of the individuals’ skills, similar to the actor’s choices. Importantly, though, whether social attention is a driver or a consequence of social evaluation (or both) and whether it is directly linked to social decisions is unclear at this point.

#### Individual and species differences in social information use

The considerable individual variation, potential species differences in performance, and the relatively small sample size may have prevented us from distilling a clearer picture. Factors like motivation, hunger, or subjects’ attention could explain individual differences. Individual features such as sex, age, personality, or rearing history can also influence the need or development of cognitive abilities and social information use. Previous studies on social evaluation (Chijiiwa et al., 2022; Hirel et al., 2024; Jim et al., 2021, 2022), social learning (Carter et al., 2014; Watson et al., 2018), or foraging activities (Jones et al., 2017; Kurvers et al., 2010; Range et al., 2009) have reported individual variation in social information use. Differences in how well individuals perform adaptive social behaviours might result from differences in their developmental history and be necessary from an evolutionary aspect by allowing behavioural flexibility to cope with changes in the social environment (Dukas, 2019; Sih et al., 2019; Taborsky & Oliveira, 2012).

The fact that the Tonkean macaques generally looked longer at the actors than the brown capuchins raises the question of species differences in social information sampling. First, the strategies for social information acquisition and use will likely vary between species with different social organisations (*e.g.*, Faraut & Fischer, 2019; Range et al., 2009; Scheid et al., 2007) and between individuals with different social status (*e.g.*, Abril-de-Abreu et al., 2015; Jones et al., 2017; Kulahci et al., 2016; van de Waal et al., 2010). For example, acquiring social information from direct experience when living in large groups can be time-consuming – interacting with each group member requires a lot of time that is not devoted to other activities. Similarly, gathering indirect social information is more beneficial than direct sampling for individuals with low tolerance or low social status, as direct interactions can be associated with a high risk of getting injured. Second, it is unclear whether potential differences in cognitive processing result in some species or individuals picking up faster than others on informational cues. Specifically, in our case, we simply do not know how much information is sufficient and required by the different monkeys to form an impression of somebody being skilful in a task.

In conclusion, Tonkean macaques and brown capuchins successfully changed their initial preference in favour of the skillful actor when this became the better option to maximise reward outcomes. They did not learn through trials to choose the better option nor did they base their choices on the outcomes experienced in previous trials. Our subjects also looked preferentially at the skilful over the unskilled actor when both actors simultaneously manipulated a container. These results indicate that these monkeys used social information about the actors’ actions acquired from the demonstration sessions to inform their decisions. Yet, whether the subjects evaluated the actors’ skills or used an outcome-based process needs to be further investigated in future studies. We encourage studies in diverse animal populations to investigate further which cognitive processes underlie social information acquisition in nonhuman animals and which social environment could favour its emergence.

## Acknowledgements

We would like to thank the management, researchers, animal keepers, and technical team of the Centre de Primatologie – Silabe de l’Université de Strasbourg (https://www.silabe.com) for their approval to conduct this study and their precious cooperation and logistical support. We are thankful to Ninon Chautard and Shadab Javanmard Paghaleh for their crucial assistance with the data collection, Nadja Vögtle for reliability coding, and Roger Mundry for the helpful advice on the statistical analyses.

## Ethics statement

This study respects the European ethical standards and regulations (Directive 2010/63/UE). It has been approved by the internal ethical committee of the Centre de Primatologie – Silabe de l’Université de Strasbourg (SBEA 2022-03, registration n° B6732636). We only used positive reinforcement and all individuals participated voluntarily – they were never forced to enter, be separated from their groups, or perform the tasks in the experimental rooms. Their daily feeding regime was not affected by cognitive testing, and water was available *ad libitum*.

## Data accessibility

The supplementary document, videos, data, and statistical code associated with this article can be found at: https://osf.io/gm7q4/ (DOI: 10.17605/OSF.IO/GM7Q4)

## Author contributions

**Hirel Marie**: Conceptualization, Data curation, Formal analysis, Investigation, Methodology, Project administration, Visualisation, Writing – original draft, Writing – review & editing. **Marziliano Michele**: Formal analysis, Investigation, Methodology, Writing – review & editing. **Meunier Hélène**: Resources, Writing – review & editing. **Rakoczy Hannes**: Conceptualization, Funding acquisition, Writing – review & editing. **Fischer Julia**: Funding acquisition, Writing – review & editing. **Keupp Stefanie**: Conceptualization, Methodology, Project administration, Supervision, Writing – review & editing.

